# CellBin:a generalist framework to process spatial omics data to cell level

**DOI:** 10.1101/2025.10.23.683357

**Authors:** Ying Zhang, Huanlin Liu, Haoxiu Wang, Zirong Li, Yumei Li, Jidong Chen, Jinghong Fan, Ji Yi, Can Shi, Xinyu Ren, Qiang Kang, Yinqi Bai, Shuangsang Fang, Jing Guo, Yang Heng, Dongmei Jia, Sha Liao, Ao Chen, Haojing Shao, Mei Li

**Affiliations:** State Key Laboratory of Genome and Multi-omics Technologies, BGI Research, Shenzhen, 518083, China; BGI Research, Shenzhen, 518083, China; Shenzhen Branch, Guangdong Laboratory for Lingnan Modern Agriculture, Genome Analysis Laboratory of the Ministry of Agriculture and Rural Affairs, Agricultural Genomics Institute at Shenzhen, Chinese Academy of Agricultural Sciences, Street, Shenzhen,518124, china; State Key Laboratory of Resource Insects, Medical Research Institute, Southwest University, Chongqing, 400715, China

**Keywords:** Spatial omics, Cell segmentation, Deep learning

## Abstract

Spatial omics has rapidly expanded with increasingly diverse imaging modalities, molecular targets, and chip sizes. However, no general framework currently exists to construct cell level matrices that are robust across platforms and omics types. Here we present CellBin, a universal and scalable framework that unifies image stitching, cell segmentation, and spot-to-cell mapping for multiple spatial omics technologies. CellBin integrates a multi-field weighted stitching algorithm for large-area images, a family of U-Net–based models trained across diverse staining modalities, and an optimized computational architecture for high-throughput processing. Across five technological platforms and three omics data types, CellBin achieves robust segmentation and accurate single-cell matrix construction, consistently outperforming seven state-of-the-art methods in F1-score, cell size precision, and annotation accuracy. By providing a generalizable, cross-platform solution, CellBin bridges multiple spatial omics, enabling unified, high-resolution cell level analyses across technologies.

## 1 Introduction

### 1.1 Introduction to spatial omics

As Method of the Year by Nature Methods in 2020[1] and 2024[2], spatial omics has advanced rapidly in recent years, with over 26 technologies already developed(Supplementary Table 1). Key features of these technologies include: 1) compatibility between spatial platforms and existing sequencing methods, enabling novel integrations such as protein CITE-seq combined with spatial platforms to create spatial-CITE-seq[3] and Stereo-CITE[4], thereby multiplying technical possibilities; 2) continuous improvements in staining methods, allowing diverse labeling of cellular components including nucleus, membrane, wall, and cytoplasm; 3) increasing chip sizes, exemplified by Open-ST[5] (6.3 × 89 mm), MAGIC-seq[6] (21.6 mm × 21.6 mm), and Stereo-Seq[7] (13.2 cm × 13.2 cm); and 4) progressively higher resolution, as seen with Visual advancing from 55 µm to 2 µm. However, the variability in data types and resolutions across platforms necessitates a unified approach for converting data into a general analysis-ready format. While single-cell analysis methods are well-established, constructing a single-cell matrix from spatial omics data offers a promising solution. Current cell segmentation methods face several limitations: 1) reliance solely on image-based segmentation without integrating spatial omics data[8, 9], 2) compatibility limited to only a few spatial omics platforms[10, 11], 3) capability restricted to low-resolution datasets with small cell numbers, and 4) support for merely one or two staining protocols[12]. To address these challenges and empower scientists to eficiently leverage spatial omics data, we developed the CellBin framework to support multiple platforms, diverse staining methods, large chips, and high-resolution single-cell matrix construction.

## 2 Results

### 2.1 CellBin model and framework

In this study, we introduced CellBin, a generalist single-cell expression matrices construction framework for spatial omics(Fig.1a). CellBin overcame existing shortcomings by developing an image stitching algorithm to enhance data preprocessing accuracy(Extended Data Fig.1e), integrating Bi-Directional ConvLSTM U-Net for nuclear images, and U-Net for cell membrane or wall images(Fig1b). Additionally, it facilitated data format conversion for Unique Molecular Identifier(UMI) of spatial single-cell[13](Fig.1c). To provide users with greater flexibility, CellBin incorporated Cellpose, U-Net for nucleus segmentation, and Gaussian Mixture Model and Euclidean Distance Map for cell segmentation, offering a scalable and versatile framework(Figure1d).

**Fig. 1.**
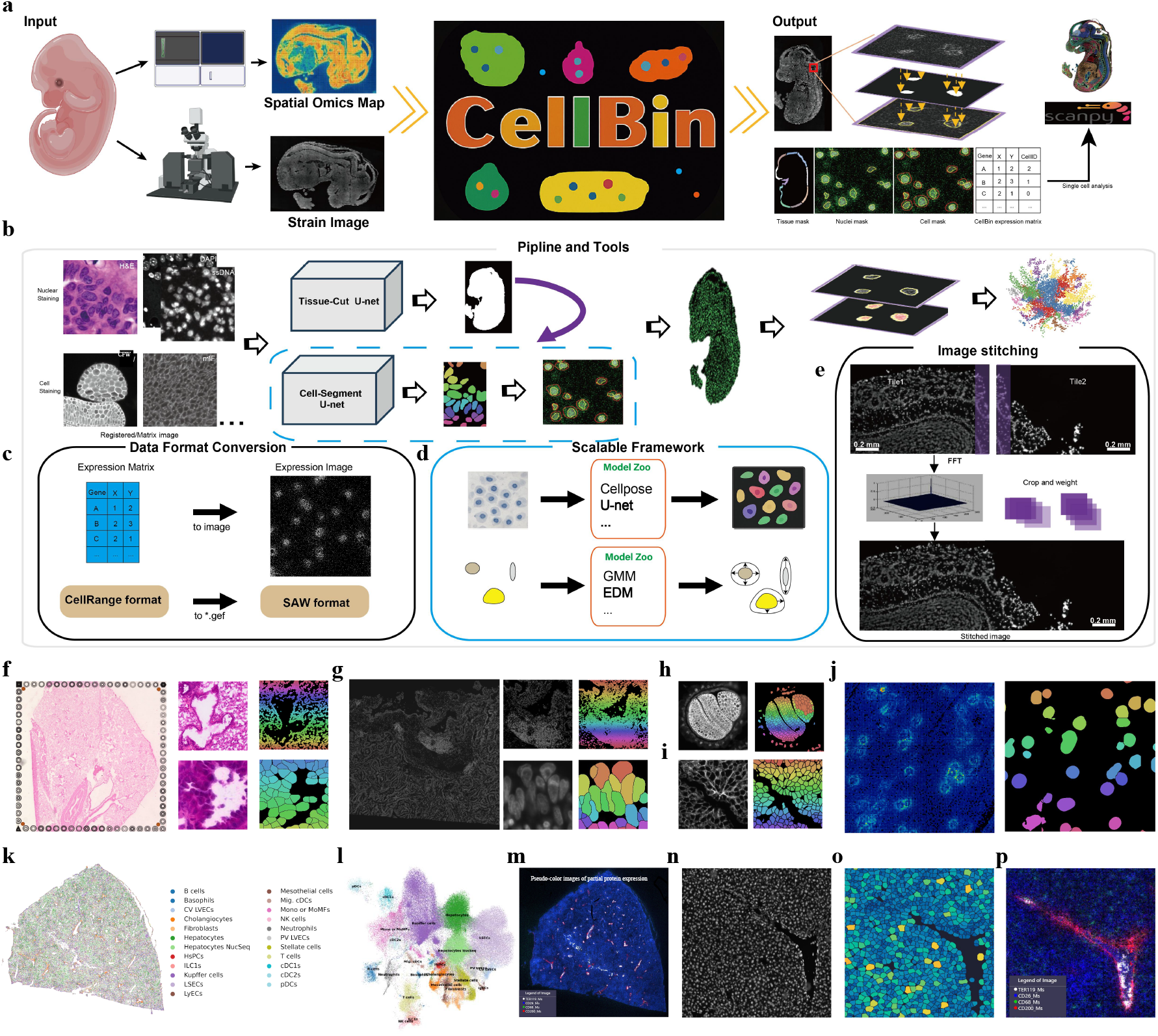
The CellBin workflow and applied examples. **a**. Schematic overview of the CellBin pipeline. The spatial gene expression data and the morphological (nuclei-stained) image of tissue are obtained using spatial transcriptomics technology and microscopy, respectively. The image tiles obtained by microscopy are stitched together to generate a large mosaic image of the whole tissue, the spatial gene expression data is transformed to a map, and the stitched image and spatial gene expression map are registered. Tissue and nuclei segmentations are performed on the registered image to obtain the tissue mask and nuclei mask. Molecule labeling is adopted to obtain single-cell spatial gene expression profile. **b**. computational overview of the pipeline; **c**, schematic of converting Stereo-cell UMI into images; **d**, scalable framework schematic; **e**, image stitching schematic; **f, g, h**, and **i** show example datasets: Visium HD H&Estained mouse lung, Xenium DAPI-stained human lung cancer, Stereo-seq CFW-stained Arabidopsis seed, and Stereo-seq mIF-stained mouse liver (left: raw images; right: CellBin results). **j** presents Stereo-cell human PBMC data (left: UMI heatmap; right: CellBin results). For the Stereo-CITE ssDNA-stained mouse liver data, **k** shows the clustering results, **i** presents the UMAP visualization, and **m** displays selected protein distributions. Zoomed-in views of a vascular region are provided in **n** (raw ssDNA image), **o** (expression heatmap after CellBin segmentation), and **p** (selected protein distribution maps).

The original staining spatial images in Spatial Omics are diverse, with mainstream staining images including DAPI, single-strand DNA(ssDNA), H&E, Calcofluor White(CFW), and Multiplex Immunofluorescence (mIF). H&E staining turns the cell nuclei purple-blue and stains the alkaline proteins in the cytoplasm and extracellular matrix red (Fig.1f). DAPI and ssDNA primarily stain the cell nucleus brightly, with the rest appearing dark(Fig.1g,n). CFW and mIF mainly stain the cell wall and cell membrane, with the cell nucleus appearing dark(Fig.1h,i,Extended Data Fig.1c). The training dataset for CellBin consisted of 4,193,768 cells across 80,907 images, covering all staining protocols(Extended Data Fig.1e), and then utilized specific U-net models to train the recognition of cell nuclei, cell membranes, or cell walls for each staining protocol individually.

Cell segmentation not only identifies cell boundaries at the pixel level but also recognizes tissue structures on a larger scale(Supplementary Fig.1). CellBin has been applied to H&E-stained 10X HD Visium mouse lung slices(Fig.1f). Lung tissue is characterized by its porous structure, comprising bronchi, alveoli, and blood vessels. These pores are visible in stained images, and cell segmentation techniques can restore their contour edges. On Xenium human lung cancer slices, multimodal staining reveals obvious tumor tissue, which CellBin can accurately identify and classify as a distinct group(Fig.1g). In Stereo-seq CFW-stained Arabidopsis seed slices, CellBin not only clearly distinguishes cell walls but also reconstructs the contours of the two cotyledons and the root stem(Fig.1h).In mIF-stained mouse liver slices, CellBin can clearly distinguishes the cell membrane (Fig.1i).

Stereo-cell[13] is a novel approach for capturing and profiling single-cell transcriptomes through spatial enhanced resolution single-cell sequencing utilizing high-density DNA nanoball (DNB)-patterned arrays. This sparse cell distribution pattern facilitates image-free, cell-specific expression profiling based solely on UMI counts. CellBin converts these UMIs into images, which are subsequently processed by U-net++ to identify cell core and boundaries(Fig.1j,Extended Data Fig.1d,2). In Figure1j and Extended Data Fig.2a, the original UMI distribution mimics ssDNA staining, with cells dispersed across the chip(Extended Data Fig.2b), indicating that UMI counts can be used to distinguish cells. The identified nucleus-like core corresponds to the region of highest expression within the cell, with signal intensity declining rapidly toward the background (Extended Data Fig.2c,d). This demonstrates that the core accurately captures the majority of the transcriptome. We further evaluated whether the cytoplasm-like ring—expanded from the core—contains the remaining transcriptome. Pearson correlation analysis reveals that the identified cell expressions closely resemble those of the core and ring(cor*>*0.71), but differ significantly from those of rings expanded by 2, 2-4, and 4-6 pixels(cor*<*0.17)(Extended Data Fig.2e). These results indicate that the ring primarily contains the remaining transcriptome, suggesting that the inferred cellular boundary is correct.

Stereo-CITE[4] is a spatial proteomics technology that integrates spatial transcriptomics Stereo-Seq with proteomics CITE-seq. CellBin performs cell segmentation, annotation, and UMAP visualization on Stereo-CITE mouse liver data(Figure1k,l, Supplementary Figure 4). The liver structure is clearly defined, with porous vascular structures distinctly visible and cell types well-classified in UMAP. Regarding captured proteins, TER119 protein (a marker for red blood cells[14]) was primarily detected in liver regions without gene expression(Fig.1m,p). Since red blood cells lack nuclei, ssDNA cannot observe nuclei or detect highly expressed DNA to identify cells(Fig.1n,o). This demonstrates CellBin’s segmentation clearly recognizes vascular voids without falsely identifying red blood cells, revealing inconsistencies between proteomics and transcriptomics. CD200 protein, expressed in vascular endothelial cells[15], was mainly captured at the edges of liver pores, spatially enveloping TER119-positive regions(Fig.1n,o). CellBin’s annotation shows cholangiocytes predominantly surround hepatic pore structures, aligning with CD200 expression patterns(Fig.1k,m). This confirms CellBin accurately delineates vascular boundaries, demonstrating consistency between proteomic and transcriptomic data. The method proves reliably applicable to both spatial transcriptomics and proteomics, correctly revealing both concordances and discrepancies between these omics layers.

### 2.2 CellBin evaluation

Processing a morphological image from a tissue slide requires the stitching of an array of multiple image tiles generated by microscopy. Microscopes can automatically capture the image tiles of a tissue one by one and stitch the image tiles together using the built-in stitching method. However, overlapping areas between adjacent image tiles generated during microscope movement may be imprecise due to mechanical tolerance and imperfect calibration. Such stitching errors are common in mosaic images and need to be removed if the goal is to achieve single-cell resolution in spatial omics applications. As an example, a mouse brain mosaic image on a chip with a size of ~2cm×2cm and a tile size of 2424pixels×2031pixels are displayed (Extended Data Fig.3a), corresponding to a physical size of ~1.2mm×1.0mm. To visualize seams and stitching errors, we select two adjacent image tiles from the image (Extended Data Fig.3b). For accurate stitching results, the lower part of tile 1 (dotted area) and the upper part of tile 2 (dotted area) should be accurately overlaid. Shadows that represent inaccurate overlap of cells can be easily seen in the stitching results produced by the microscope (Extended Data Fig.3c, left), but MFWS of CellBin is able to stitch the image tiles accurately, resulting in the absence of shadows (Extended Data Fig.3c, right). The stitching results of two image tiles within non-tissue areas (full-colored part) where individual “track lines” can be clearly seen using auto-contrast adjustment as also shown (Figure2a). The applied Stereo-seq chip contains periodic “track lines” with a size of 1500 nm (~3 pixels), which is apparent from the image (Extended Data Fig.3c).

We use a collection of data to evaluate the accuracy and efficiency of existing stitching algorithms as compared with MFWS. We apply MFWS to the public dataset, and the results show that the relative offset errors of MFWS are comparable with those of MIST[16], both being concentrated within 5 pixels, while ASHLAR[17] has larger offsets (*>*10-pixels) (Fig.2b). Moreover, the offset error distribution is much more concentrated by MFWS. Next, we calculate the relative and absolute offset errors on four gradually increasing stereo-seq samples (ranging from 1cm×1cm to 2cm×3cm, Fig.2c). MFWS is shown to perform significantly better than ASHLAR[17] and MIST[16] with respect to the relative offset errors for all image size combinations (Extended Data Fig.3d). A 10-pixel (~5 m) dislocation roughly corresponds to half a cell (Fig.2a). In all the datasets, the errors generated by MFWS remain within a maximum of 10-pixels in the tissue area, while the other methods produce a shift greater than 10-pixels, which may misplace the cell (Fig.2c). As the number of image tiles increases, the run time becomes a significant factor for stitching algorithms. The run time of MFWS is significantly shorter than that of the other methods, mostly due to the embedded spectral calculation based on FFT (Fig.2d).

**Fig. 2.**
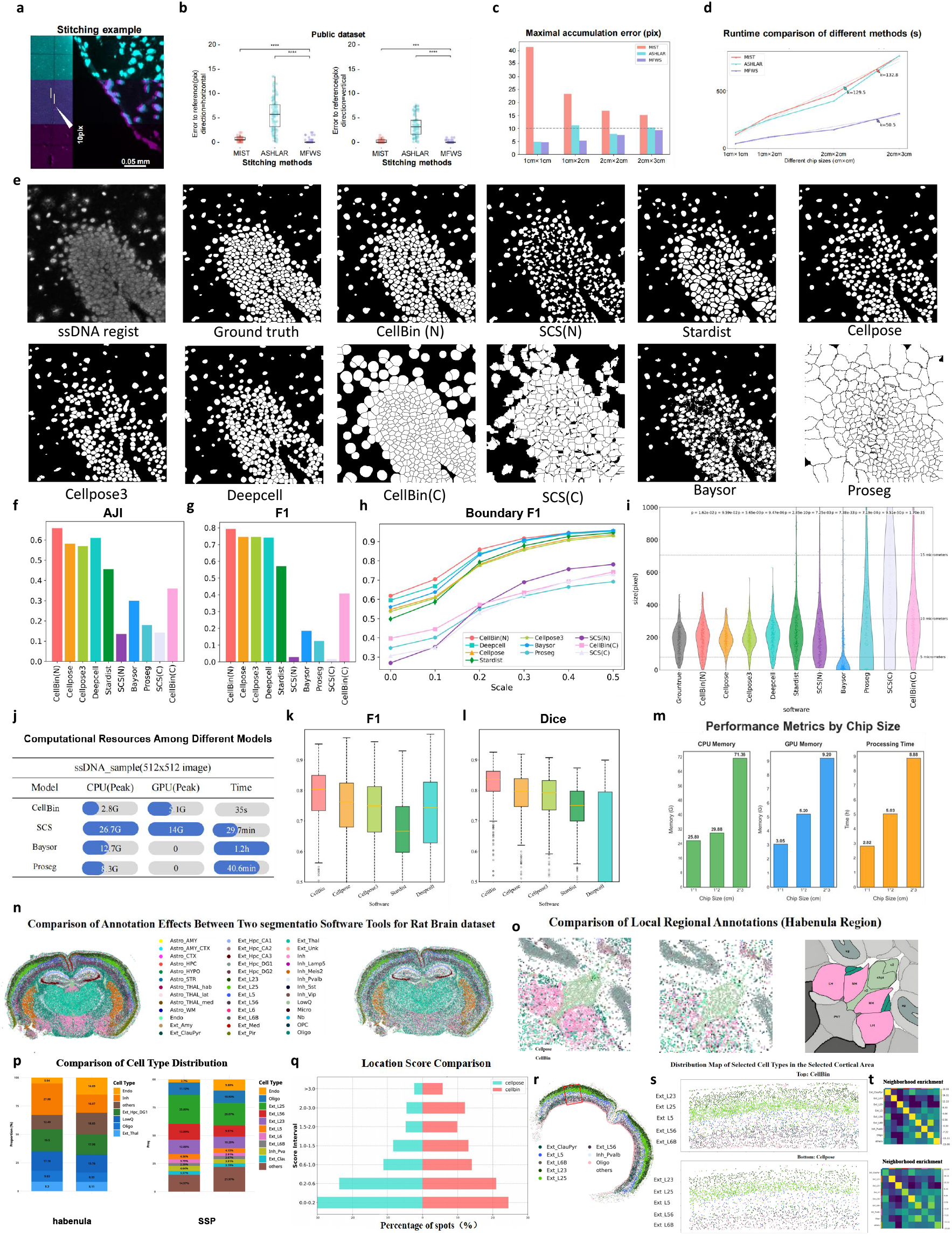
The Evaluation of CellBin. **a**. An example of the stitching error in pixels in the zoomed-in image. A dislocation of 10-pixels roughly corresponds to half a cell. **b**. Comparison of the relative errors produced by MIST, ASHLAR, and MFWS on a public dataset.**c**. Bar graph illustrating the maximal accumulation of stitching errors produced by MIST, ASHLAR, and MFWS on the datasets. **d**. Line graph displaying the run time of MIST, ASHLAR, and MFWS on Stereo-seq mouse brain datasets. **e** are 512×512 pixel examples of ssDNA stained data. The first images depict raw data, the second show manually annotated ground truths, and subsequent images exhibit cell segmentation results from various methods. For ssDNA examples, **f, g, h, i, j** represent AJI (Aggregated Jaccard Index), F1 score, boundary F1, cell size, and computational resource respectively. **k** and **l** display the F1 and Dice scores for 482 image sets[18]. **m** illustrates CellBin’s CPU memory usage, GPU memory consumption, and processing time across chip sizes of 1cm×1cm, 1cm×2cm, and 2cm×3cm. **n** shows the cell2location annotation results for complete mouse brain slices using CellBin and Cellpose. **o** left and middle present magnified views of the third brain region for CellBin and Cellpose, while **o** right depicts the corresponding region in the Allen Mouse Brain Atlas. **p** compares the cell proportions in two distinct regions. **q** displays the location scores for cell2location across the entire mouse brain using both tools. **r** visualizes the cerebral cortex of the entire mouse brain processed by CellBin. **s** and **t** provide magnified views of the SSp cortex and its neighbor analysis z-scores, respectively.

Cellbin is compared with four pure image-based cell segmentation methods: Cellpose, Cellpose3, Deep-cell, and Stardist, as well as three ST “nucleus-expansion” based cell segmentation software: SCS, Baysor and Proseg. The test sets include Stereo-Seq ssDNA-stained mouse brain data (Fig.2e) and H&E-stained mouse liver data(Extended Data Fig.4). Two 512*512 pixel regions, each containing 397 and 517 cells, are extracted. True masks are determined by ground truth IOU*>*0.5 to calculate TP, FP, and FN. Overall accuracy is quantified by AJI(Fig.2f) and F1(Fig.2g). For cells matching the ground truth, boundary F1(Fig.2h) and cell size (Fig.2i) are used for comparison. To demonstrate the universality of this superiority, we compared CellBin with other software using 482 imaging datasets[18](Supplementary Figure 6). CellBin consistently outperformed all alternatives(Fig.2k,l).

CellBin substantially outperforms other software across all quantifiable metrics. In ssDNA staining data, CellBin, Cellpose, Cellpose3, DeepCell, and Stardist are relatively close to Ground Truth, with Cell-Bin being the closest(Fig.2f). Cellpose, Cellpose3, DeepCell, and Stardist miss some cells in dense areas, while SCS and Baysor miss even more. In H&E staining data, CellBin’s recall is 0.75, compared to Stardist’s 0.39, which ranks second(Extended Data Fig.4k). Overall, CellBin also excels in F1 and AJI metrics. When comparing cell sizes to Ground Truth, all software tools exhibited substantial discrepancies—Baysor tended to detect excessively small cells while Proseg identified disproportionately large cells, whereas only CellBin closely matched the Ground Truth size measurements(Fig.2i). For a more detailed comparison, we expanded cell boundaries from 0 to 0.5 cell radii, and CellBin outperformed other software across all expansion sizes(Figure2h).

Notably, CellBin demonstrates superior image generalization capability. H&E staining presents a challenging scenario where nuclei appear blue, cytoplasm stains purple, and interstitial regions remain white. SCS is hard to process color images, while DeepCell merges cells in purple-stained cytoplasmic regions, leading to under-segmentation.

Subsequently, a comparison of downstream cellular analysis was conducted between CellBin and Cellpose in this mouse brain tissue(Fig2n). CellBin detected 44% more cells overall than Cellpose and 257% more transcriptomes. The average number of transcriptomes per cell was 356 for CellBin and 144 for Cellpose. Using cell2location[19] for annotation of the mouse brain, CellBin achieved a 19% higher annotation score than Cellpose(Fig2q). As a result, CellBin demonstrated higher cell density and clearer overall annotation. Taking the third ventricle as an example, CellBin clearly identified the habenula region, whereas Cellpose’s habenula was less distinct and mixed with other cells (Fig2o,Supplementary Figure 3). Further analyzing cell proportions, CellBin had 11.9% more lnh cells than Cellpose, while Cellpose had 9.9% more Endo cells than CellBin(Fig2s). Analysis of Endo cells reveals that across the entire slice, Cellpose’s Endo cells are more diffusely distributed than CellBin’s(Supplementary Figure 2), suggesting that inaccurate cell segmentation may increase the apparent proportion of Endo cells. Similarly, in the primary somatosensory cortex (SSp), both CellBin and Cellpose detect the cortical layer structure of SSp, with CellBin exhibiting denser and clearer cell patterns(Fig.2s). Regarding cell proportion, Cellpose has 7.2% more Endo cells than CellBin(Fig.2p). Subsequent neighbor analysis reveals that CellBin achieves higher z-scores in self-comparison across cell types compared to Cellpose (Fig.2t), indicating that CellBin’s cell organization contains fewer misclassified cells. Consequently, when CellBin detected more and more accurate transcriptomes, cell2location’s annotation became more precise, reducing the likelihood of cells being misidentified as others and ensuring more accurate cell proportions.

CellBin exhibits significant advantages over other ST cell segmentation software in terms of speed and memory usage. In 512×512 pixels screenshots, CellBin is at least 50 times faster than SCS, Baysor, and Proseg(Fig.2j). The CPU memory consumption of SCS, Baysor, and Proseg is at least 1.6 times that of CellBin, while SCS’s GPU memory usage is 2.74 times that of CellBin. In complete slices with 23520 * 23520 pixels, where the number of cells increases by 50-100 times, CellBin can still complete the task in approximately 1 hour(Extended Data Fig.5).

The expansion of spatial omics chips represents a key advancement in this technology[5, 6]. CellBin can also be applied to large-scale chips(Supplementary Figure 5). CellBin’s MFWS algorithm ensures stitching accuracy for large chips, while its high computational efficiency enables feasible processing of such large-scale data. On a 2cm×3cm chip, CellBin requires only 72GB of memory, 10GB of GPU memory, and 9 hours of CPU computation(Fig.2m).

### 3 Short summary

CellBin constructs single-cell matrices across image-based and sequencing-based spatial transcriptomics, spatial proteomics, and recently emerging spatial single-cell datasets[13]. It incorporates multiple models tailored for diverse spatial omics image data, including ssDNA, DAPI, mIF, CFW, H&E, and even UMI-based images. In multi-dimensional benchmarks against seven state-of-the-art tools, CellBin consistently outperforms competitors. It also demonstrates superior clarity and reliability in downstream cell annotation analysis. Additionally, CellBin’s independently developed image-stitching algorithm surpasses existing methods, ensuring high accuracy for larger spatial omics chips. It also exhibits computational efficiency and resource optimization, enabling scalability for extensive datasets. In summary, CellBin serves as a universal framework to process spatial omics data at the single-cell level, well-suited for the rapidly evolving field of spatial omics technologies.

## 4 Method

### 4.1 Image stitching

The image tiles are stitched into a large mosaic image of the whole tissue by MFWS (Fig.1e and Extended Data Fig.1a). In contrast to conventional microscope techniques dependent on hardware-based overlap parameter, MFWS utilizes cross-power spectral analysis to achieve higher-precision overlap detection through similarity maximization. We assume that *a*(*x, y*)?and *b*(*x, y*)?are two tiles need to be stitched, *x* and *y* represent coordinates. Based on the microscope’s hardware overlap parameters, their overlapping region can be extracted as *a*^*′*^ and *b*^*′*^. Firstly, *a*^*′*^, *b*^*′*^ is divided into *n* smaller sub-images through resampling: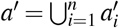and 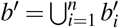Then apply the Sobel operator to filter to remove sub-images with insufficient features, resulting in *m* qualified sub-images. We can compute the Fourier transforms of 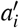 and 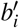, and then calculate their cross-power spectrum. Fourier Transforms as follow:

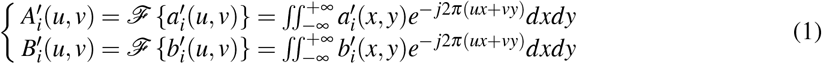

ℱ{·} is Fourier transform operator.

Two sub-images normalized Cross-Power Spectrum *C* is:

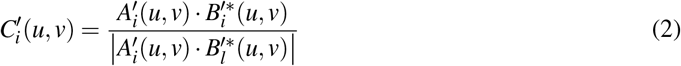

Where * means complex conjugate operator.

Due to the low signal-to-noise ratio of some sub-images in the previous step, the calculated value deviates significantly from the actual value. The overall accuracy of the algorithm can be significantly improved by applying a weighted enhancement to partial blocks and a reduction to residual sub-blocks. Our weighting coefficient is based on the peak value of the cross power spectrum because experiments show that the greater the peak value, the higher the precision and reliability of the the estimated overlap offset. A unique cross-power spectrum is obtained after the weighted superposition of *m* spectra. Then, the corresponding spatial domain correlation graph can be obtained through inverse fast Fourier transform(IFFT). The coordinate of the peak value represents the required offset value. The cross-power spectra for overlap calculation between the two final tiles are shown below:

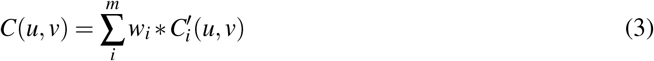

Where *w*_*i*_ is weight matrix, defined as: 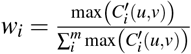

From the superimposed cross-power spectrum *C*(*u, v*), we can obtain the position corresponding to the maximum response in the global matching score matrix. This position is identified as the optimal stitching overlap offsets.

We acquire image pairs by scanning the image rows and columns. Our algorithm then iterates through all candidate pairs to calculate their optimized overlap offsets, ultimately reconstructing the complete mosaic image.

### 4.2 Image registration

Spatial omics technologies utilize distinct registration methodologies depending on their respective platforms. For instance, the 10X Visium HD platform employs fiducial markers encircling the chip as reference points for spatial alignment. CellBin recommends that users adhere to the standardized registration protocol provided by the manufacturer and input pre-aligned images into the pipeline. Furthermore, we have implemented a marker-based registration algorithm specifically optimized for Stereo-seq data (Extended Data Fig.1b).

Image registration was performed based on the fiducial markers pre-fabricated on the Stereo-seq chip. These markers were automatically detected in both the high-resolution tissue image and the sequencing coordinate system derived from the expression matrix. The registration process consisted of a two-stage approach: First, a coarse alignment was achieved by translating the feature image to minimize the distance between the global centroids of the marker points from both sources. Following this initial transformation, a fine, rigid refinement was applied. This step involved optimizing the spatial transformation (translation and rotation) to precisely register the full set of corresponding marker points, maximizing their pairwise correlation. This hierarchical strategy ensured robust and high-precision alignment, which is critical for accurate image registration.

### 4.3 Tissue & Cell segmentation

Model: For tissue segmentation, we employed two specialized Bi-Directional ConvLSTM U-Net models [20](Extended Data Fig.1c), each tailored to a specific image type: one optimized for single-channel images and the other for H&E-stained samples. This dual-model design enabled effective capture of spatial and contextual features within heterogeneous histological data. For nuclear segmentation, we extended the same architectural framework by developing two additional models, also based on the Bi-Directional ConvLSTM U-Net, which were specifically optimized for single-channel and H&E-stained nuclear images, respectively. To segment cellular membranes (MIF-stained) and cell walls (CFW strain), we first applied an inverted color transformation to improve structural contrast, followed by segmentation using the Cellpose framework, which utilizes pre-trained models for robust and generalized cellular boundary detection [18].

Training: The model is trained by minimizing the loss function *ℒ*, which is binary cross-entropy:

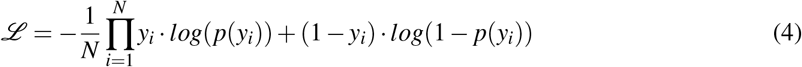

Where *y* is a binary label, either 0 or 1, and *p*(*y*)?represents the probability of the output being *y*. The binary cross-entropy serves as a loss function to evaluate the performance of a classification model.

We developed a dual-path segmentation framework comprising two specialized weight models optimized for single-channel grayscale and H&E-stained images respectively, trained on an extensive dataset of 1632 manual annotated Frozen Fresh tissue samples (Extended Data Fig.1e). For nuclear segmentation, we developed two specialized deep learning models trained on distinct imaging modalities: ssDNA (including DAPI-stained), H&E images, leveraging a large-scale dataset containing tens of thousands of images with approximately 4 million expertly annotated nuclear masks (Extended Data Fig.1e).The annotations for cell segmentation were generated using a semi-automated approach. An initial model was trained on a small set of manually annotated images and was used to propose segmentation masks for a larger image pool. All proposed masks were then meticulously inspected and manually corrected by trained experts to ensure maximum accuracy for model training.

Preprocessing and postprocessing pipelines were specifically designed for each computational model to optimize their performance on distinct histological image types and analytical tasks. The detailed procedures are summarized in Supplementary Table 4.

To enhance the quality of cell-level expression matrices, we implemented a spatial filtering strategy by computing the intersection between tissue segmentation masks and cell/nuclear segmentation results.

### 4.4 Stereo-cell segmentation

Stereo-cell segmentation is performed without requiring a tissue segmentation step. First, the spatial gene expression matrix is transformed into a grayscale image where the value of each pixel represents the total UMI count. The grayscale histogram of this image is then computed to identify the valid intensity range [*h*_*min*_, *h*_*max*_]. Contrast stretching is applied by clamping pixel values below *h*_*min*_ to *h*_*min*_ and values above *h*_*max*_ to *h*_*max*_. The remaining values within this range are linearly normalized to enhance contrast.

Image segmentation is performed using a U-Net++ model[21] (Extended Data Fig.1d). Pixel values after sigmoid activation greater than 0.5 are assigned to the foreground, and the rest to the background. Small objects are filtered out, and the watershed algorithm is applied to address cell adhesion issues.Finally, instance segmentation results are converted into a semantic representation, producing a binary mask that can be directly utilized to extract single-cell expression matrices.

The U-Net++ segmentation model was trained on a dataset of 6,070 manually annotated cells. The Focal Loss is:

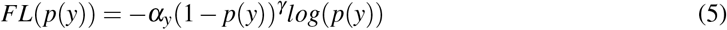

Where *p*(*y*)?is the the model’s estimated probability for true class, same shown in equation (4); *α*_*y*_ is the class balancing factor for the true class *y. γ* is the focusing parameter that adjusts the rate at which easy examples are down-weighted.

### 4.5 A nuclear-to-cellular expansion

CellBin directly extracts single-cell expression matrices from cellular segmentation masks derived from cell membrane/wall staining. However, when processing nuclear masks, the resulting matrices inherently capture only intra-nuclear expression profiles. To address this limitation, CellBin provides two nuclear-to-cellular extrapolation methods: EDM Rapid Dilation (Extended Data Fig.1f) and GMM-based method.

The EDM algorithm assigns spatial transcriptomics spots to cells through a two-step process: first, it excludes spots with Euclidean distances (computed via distance transform) that exceed a predefined threshold *Z*; second, it allocates the remaining spots to their nearest neighboring cells based on a minimum distance metric.

GMM is used to model the molecular distribution surrounding each cell. To develop the cell mask, we first expand the nuclear mask boundary to capture as much of the true cellular data distribution as possible, rather than just the nuclear distribution. Based on empirical data regarding cell area, we set the fitting range to a fixed size of 100 pixels × 100 pixels. Subsequently, the GMM calculates a probability score for each extracellular molecule within this fitting range. Finally, an adaptive threshold for the current cell is determined from these probability scores using the interquartile range (IQR). Molecules with a probability score greater than this threshold are reassigned to the cell. This approach ensures that molecules are assigned to their corresponding cells with high confidence.

### 4.6 Data format conversion

The CellBin pipeline utilizes the GEF and GEM formats (generated by Stereo-seq) as the default for storing processed expression matrices. These formats is compressed and incorporates spatial coordinates along with gene expression counts. The resulting single-cell expression matrix is also saved in the GEF format. Both GEF and GEM files can be conveniently converted into more commonly used downstream analysis formats, such as AnnData or Seurat objects, using the Stereopy package [22]. Additionally, CellBin provides a utility for converting 10X Genomics output data into the GEF format. Data in other custom formats can be similarly transformed by following an analogous conversion procedure.

## Acknowledgements

We thank the China National GeneBank for providing data storage support for this study and. This work was supported by the National Key R&D Program of China (2022YFC3400400) and the National Natural Science Foundation of China (Grant No. 32200517). We thank Zhonghan Deng, Yufeng Deng, for their assistance.

## 5 Declarations

Some journals require declarations to be submitted in a standardised format. Please check the Instructions for Authors of the journal to which you are submitting to see if you need to complete this section. If yes, your manuscript must contain the following sections under the heading ‘Declarations’:

### Data availability

A summary of all involved datasets is given in Supplementary Table 5 The raw sequencing data has been deposited into CNGB Sequence Archive (CNSA) of China National GeneBank DataBase (CNGBdb) with accession number CNP0007817: https://db.cngb.org/search/project/CNP0007817; PBMC dataset is archived at https://db.cngb.org/dataresources/sample/CNS1034433; Human Lung cancer of Cancer: https://www.10xgenomics.com/datasets/preview-data-ffpe-human-lung-cancer-with-xenium-multimodal-cell-segmentation-1-standard;

Mouse Lung of Visium HD:https://www.10xgenomics.com/datasets/visium-hd-cytassist-gene-expression-mouse-lung-fresh-frozen;mIF-stained Mouse Live and CFW-stained Arabidopsis seed:https://github.com/STOmics/STCellbin.

### Materials availability

### Code availability

Github project homepage: https://github.com/STOmics/cellbin2/tree/paper

Programming language: Python

Other requirements: Python 3.8

License: MIT License

If any of the sections are not relevant to your manuscript, please include the heading and write ‘Not applicable’ for that section.

---

**Extended Data Fig. 1.**
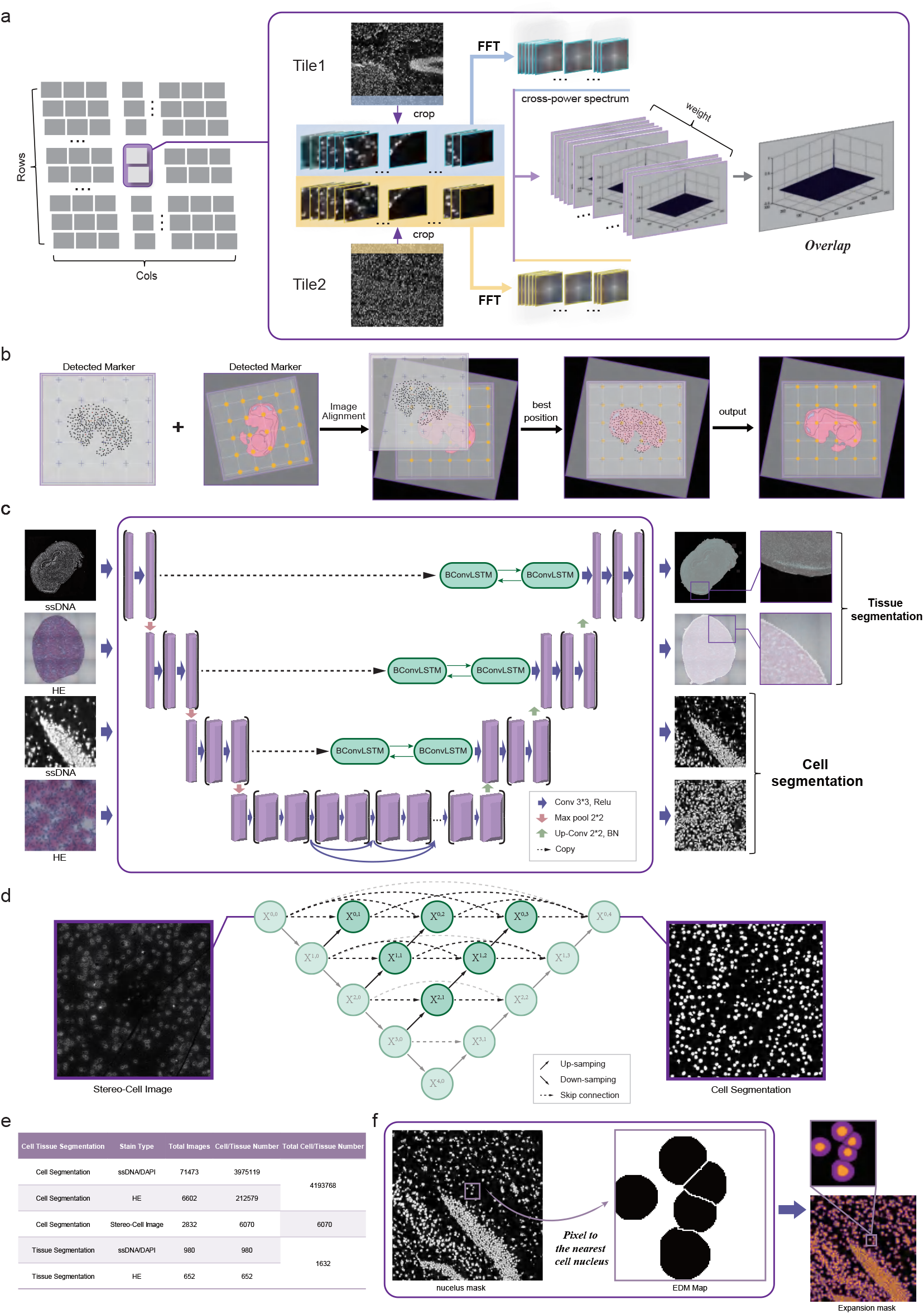
Schematic illustration of the underlying algorithm employed by CellBin. **a** Image stitching: the left images represents the microscope tiles to be stitched together in an experiment. Taking two of the tiles as an example, their overlapping regions are extracted and divided into multiple small sub-images. Some sub-images with rich features are selected to compute the cross-power spectrum. A more accurate overlap value is then generated by weighting the multiple cross-power spectrum results. **b** Image Registration for Stereo-seq: The intersection points of the track lines can be detected in the transformed map and stitched image by line searching algorithms. Then image-to-matrix registration can be optimized using a set of acquired points. Applying the estimated transformation parameters to the image yielded a registered image. **c** 4 Models of cell and tissue segmentation [20] **d** CellBin utilizes a U-Net++ architecture for cell instance segmentation on Stereo-cell[13] images. **e** The training set size for cell and tissue segmentation models **f** Introduction to the EDM algorithmic workflow and demonstration of results.

**Extended Data Fig. 2.**
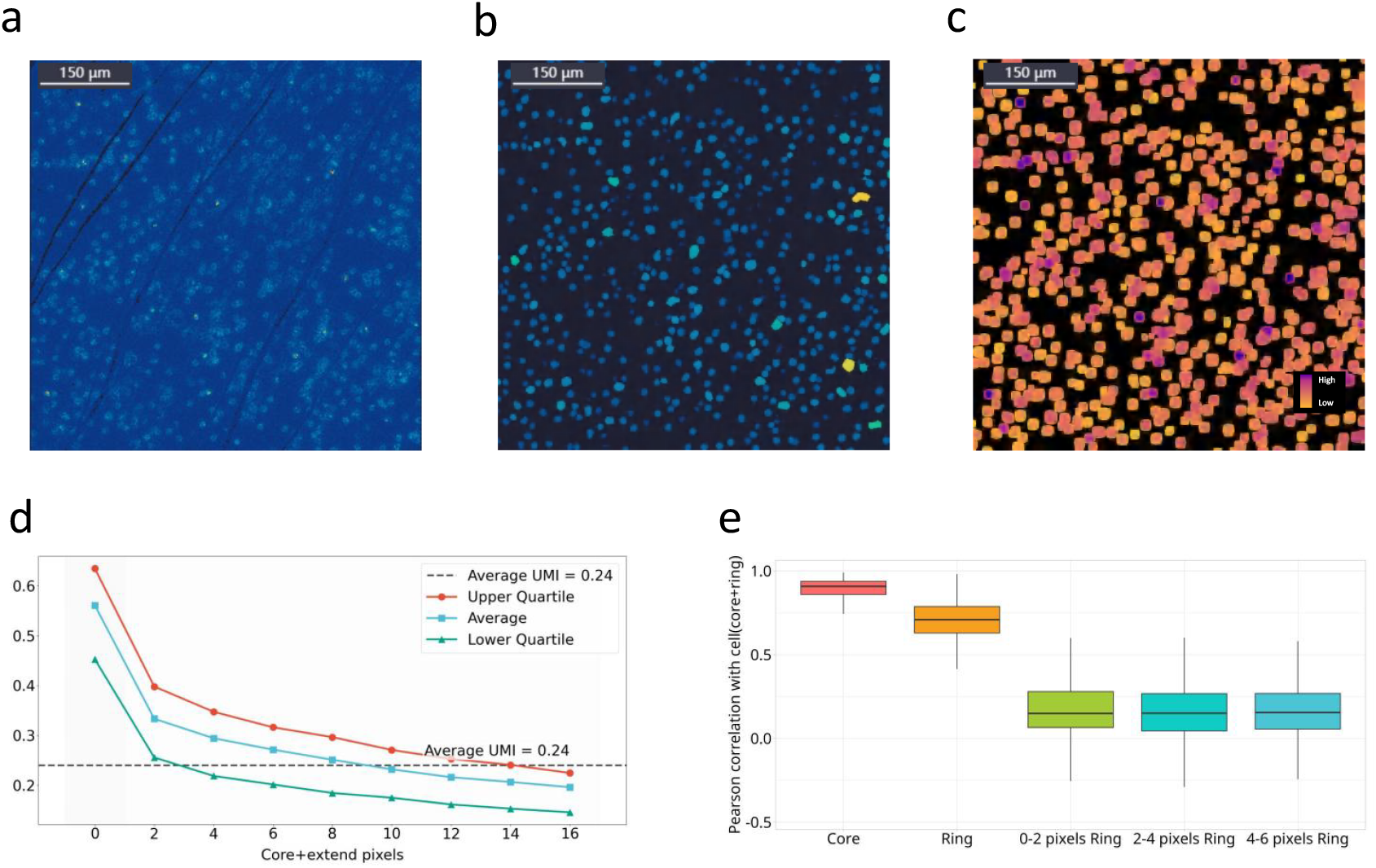
Examples and evaluation on Stereo-cell UMI-only human PBMC datasets. **a**. UMI heatmaps visualizing single-cell transcriptome data. **b**. CellBin results displaying cell classification. **c**. average UMI extension from CellBin-identified UMI cores. **d**. average UMI distribution from the core (0 pixel) to 16 extended pixels, incremented by 2 pixels each time. **e**. Pearson correlation analyzing overall gene expression between cells (nucleus-like core+cytoplasm-like ring) and five regions—core, ring, 0-2 pixels ring, 2-4 pixels ring, 4-6 pixels ring. X-Y pixels ring refers to areas extending from X pixels to Y pixels from the cell core.

**Extended Data Fig. 3.**
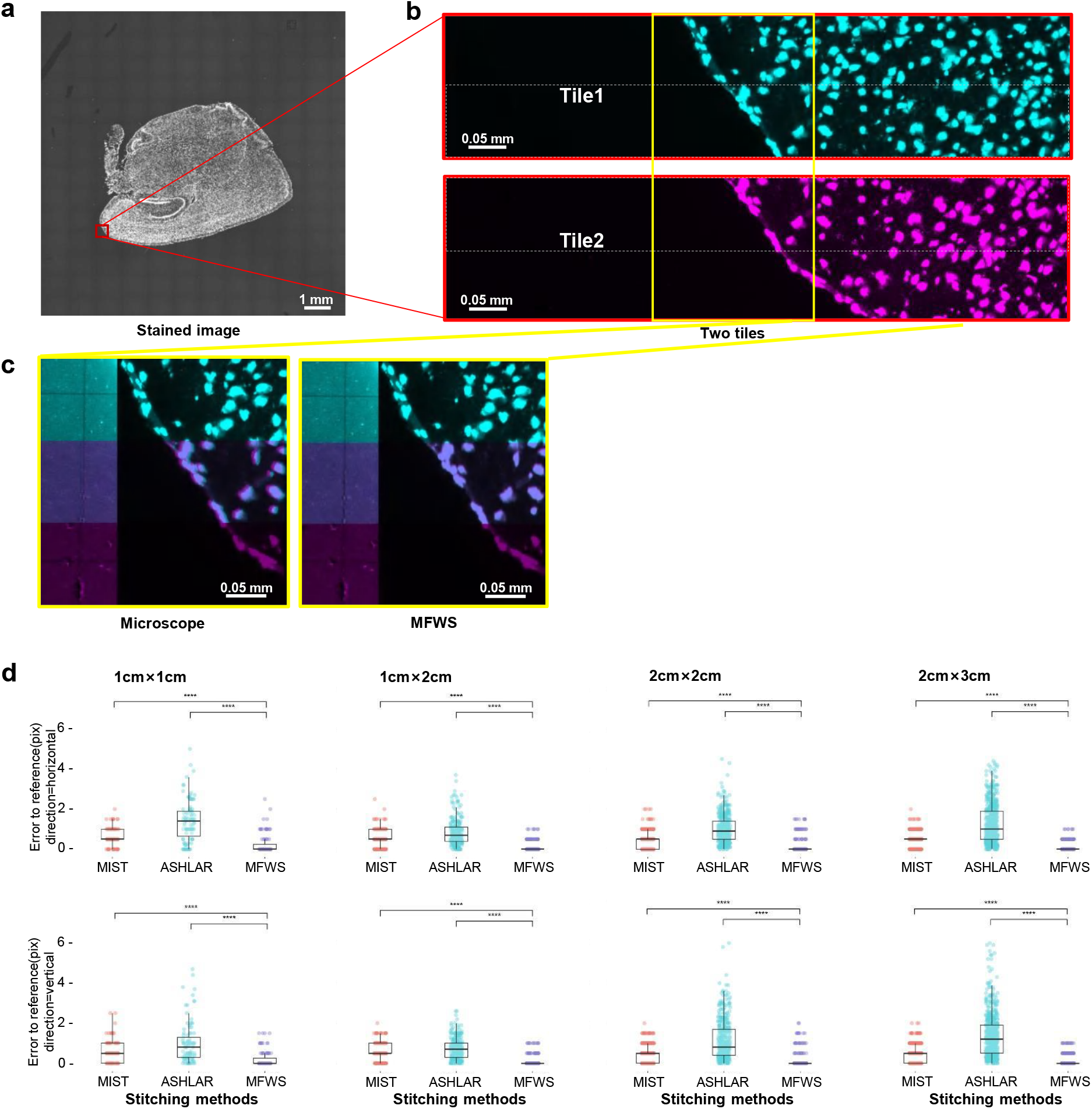
CellBin includes a high-precision mosaic method for image stitching, enabling large field-of-view morphological image analysis at single-cell spatial accuracy. **a**. Mosaic image of mouse brain tissue on a 2cm×2cm chip. **b**. The red box indicates the edge of the tissue (containing cells and non-tissue), and two neighboring image tiles (in the vertical direction) are shown in different colors. The part that needs to be overlaid by stitching is the lower part of tile 1 (within the dotted area) and the upper part of tile 2 (within the dotted area). **c**. The stitching results for the two image tiles (yellow box in b) using the stitching method built into the microscope (left) and MFWS (right). In each sub-figure, the left part shows the overlap of cells after stitching and the right part shows the “track lines” in the non-tissue area. The contrast of the background was increased to clearly show the “track lines” incorporated into the Stereo-seq chip. **d**. Comparison of the relative errors produced by MIST, ASHLAR, and MFWS on Stereo-seq mouse brain datasets, analyzed using different chip sizes from 1cm×1cm to 2cm×3cm, corresponding to number of image tiles from 11×9 to 23×29. The evaluation is based on the ground truth calculated using the “track lines” on the Stereo-seq chip.

**Extended Data Fig. 4.**
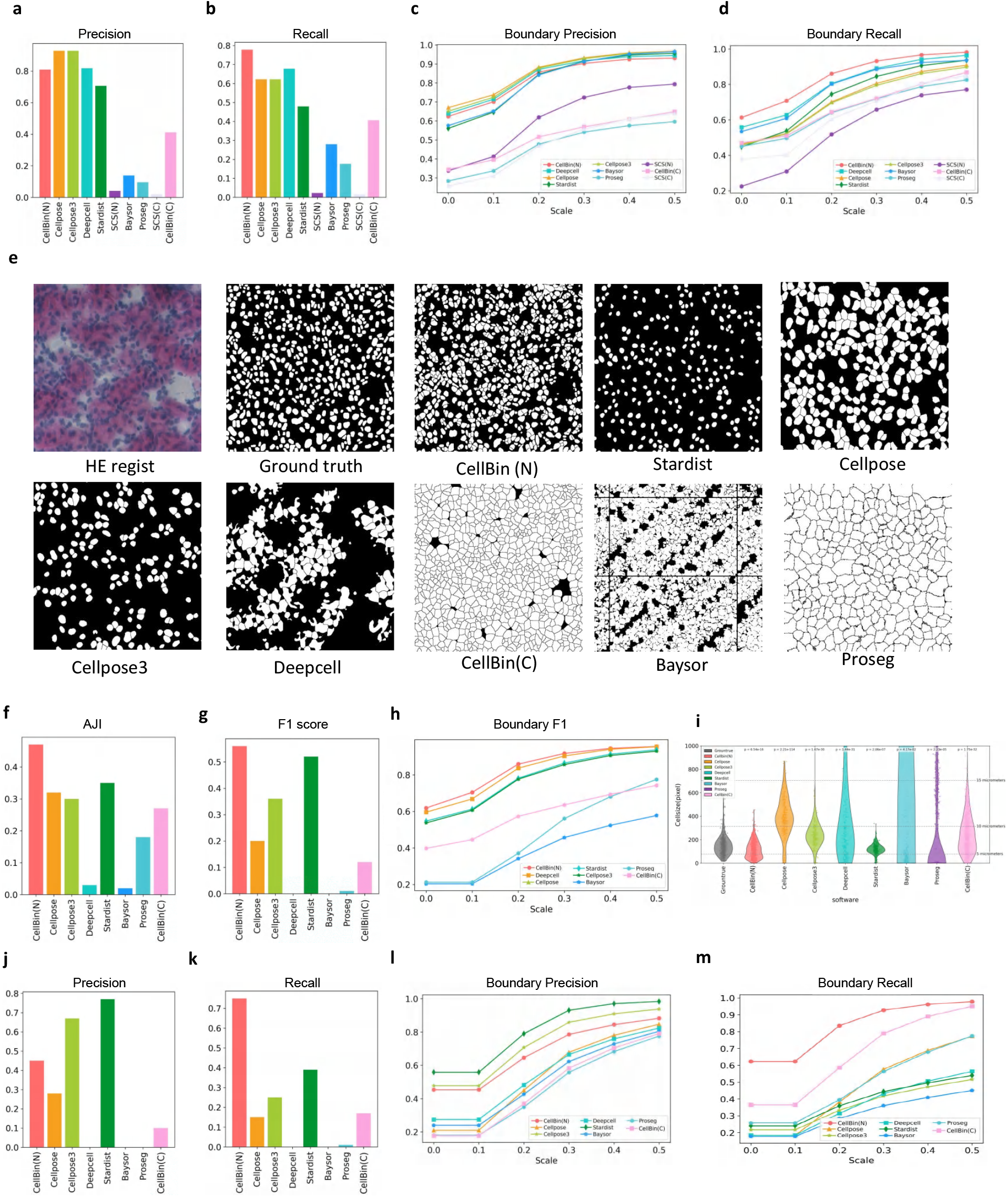
The supplementary for Evaluation of CellBin. **a** and **b** indicate precision and recall for 512*512 ssDNA images; **c** and **d** show precision and recall of boundary scores for ssDNA images; **e**are 512×512 pixel examples of HE stained data.The first images depict raw data, the second show manually annotated ground truths, and subsequent images exhibit cell segmentation results from various methods; For HE exmaples, **f,g,h,i,j,k,l,m** represent AJI, F1 score, boundary F1, cell size,precision,recall,boundary precision,boundary recall respectively.

**Extended Data Fig. 5.**
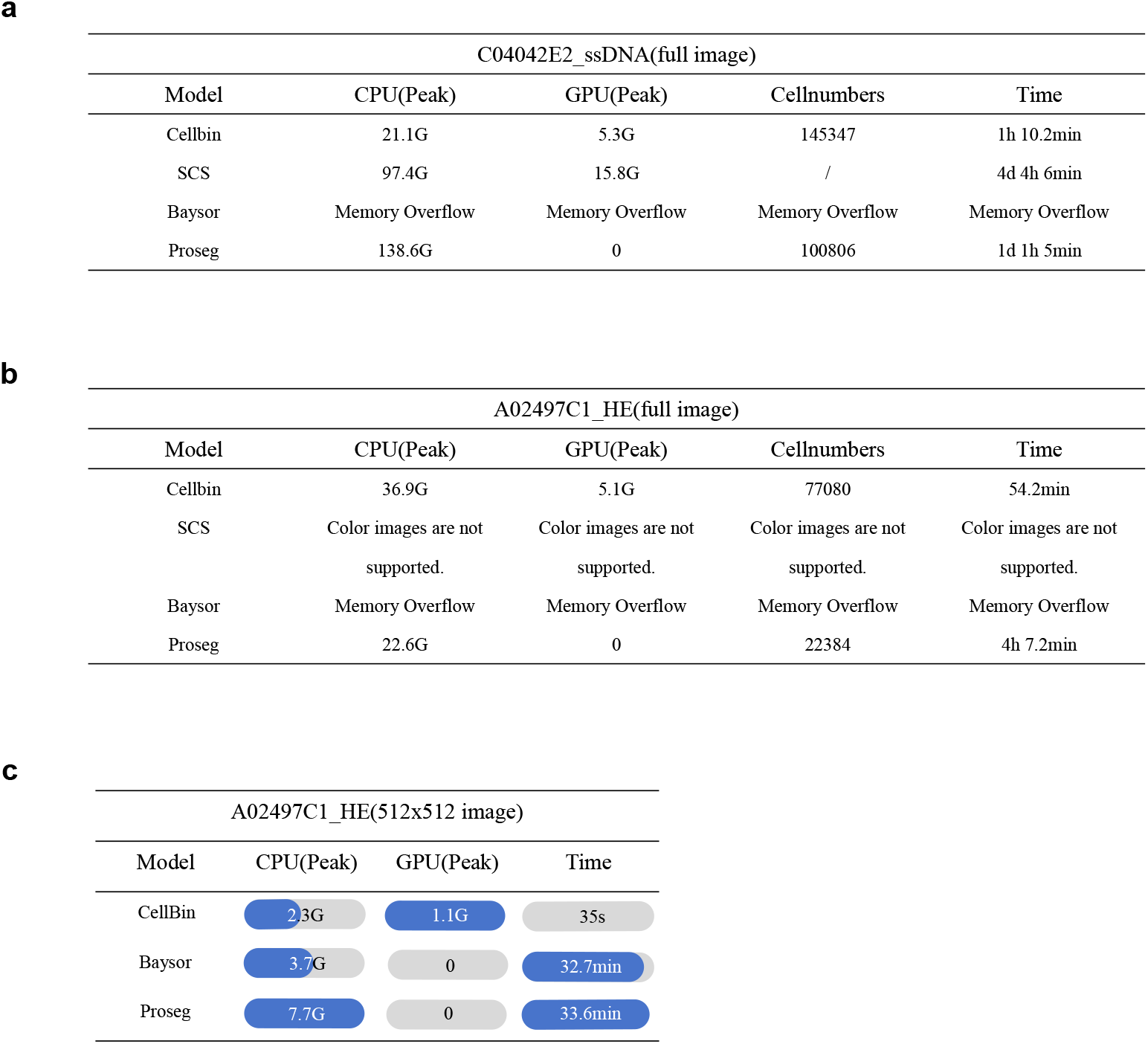
Computational resource comparison of CellBin, SCS, Baysor, and Proseg on complete datasets. **a** for full image ssDNA sample; **b** for full image H&E mouse sample; **c** for 512×512 H&E mouse sample.

