## supplementary for "CellBin:a generalist framework to process spatial omics data to cell level"

### SupplementaryNotes

#### Supplementary Note 1:Xenium and Visium HD data preprocessing

Xenium human lung cancer and Visium HD mouse lung tissue datasets including the gene expression matrices and paired high-resolution microscopy staining images, were obtained from the 10X Genomics official website. The strain images were pre-aligned to the expression data already. CellBin provide preprocessing step to convert *.h5 into the *.gef.  ubsequently, the converted GEF file and its paired microscopy image were used as the direct input for the CellbinV2 analytical suite.

#### Supplementary Note 2:Cell type annotation

We performed cell type annotation on mouse brain tissue using cell2location with the reference dataset provided directly by the cell2location (V0.1.4) package. For mouse liver data, cell annotation was conducted via label transfer implemented in the Seurat toolkit. The expression matrix was first normalized and log-transformed, followed by dimensionality reduction. Anchors between the reference and query datasets were identified using the Seurat Integration method, after which label transfer was applied to complete the annotation.

#### Supplementary Note 3:Evaluating the Performance with molecular labeling method

We compared CellBin with several established molecular labeling algorithms, including Baysor, SCS, and ProSeg. Baysor and ProSeg utilized the cell segmentation results generated by the officially recommended Cellpose algorithm as input. SCS employed its own nuclear segmentation method to generate seed points for cell demarcation. For the comparative evaluation of time consumption, memory usage, and GPU memory efficiency, only the procedures involving cell segmentation, Euclidean Distance Map (EDM) calculation, and expression matrix extraction were considered in the analysis of CellBin.  CPU and GPU telemetry were profiled on an NVIDIA T4 (CUDA 11.7) system using psutil (v7.0.0) and pynvml (v12.0.0), respectively.

#### Supplementary Note 4:Computational Efficiency on large-scale chips

To rigorously evaluate the scalability and computational efficiency of CellBin in processing large-scale tissue chips, we performed a comprehensive benchmarking analysis. Tissue samples of increasing sizes (1X1 cm, 1X2 cm, and 2X3 cm) were processed through the complete CellBin pipeline, which includes image registration, tissue segmentation, cell segmentation, and single-cell matrix extraction. The execution time for the entire workflow was meticulously recorded. Furthermore, system resource consumption was monitored programmatically: CPU utilization was tracked using the psutil library (v7.0.0) to measure computational load, while GPU telemetry was collected programmatically using pynvml（V12.0.0）on Linux system equipped with NVIDIA T4 GPU, CUDA 11.7.

#### Supplementary Note 5:STOmics Data Collection

A subset of the data presented in this study was generated using Stereo-seq and Stereo-CITE-seq technologies, with all sequencing data deposited in the CNGBdb database. Tissue samples were harvested from 5-week-old male C57BL/6J mice and 10-week-old male Sprague-Dawley (SD) rats, and all animal-related experimental procedures were performed in strict compliance with ethical guidelines for animal research.

Fresh-frozen tissue samples were sectioned at a 10 μm thickness following the STOmics Transcriptomics Set User Manual, and sections of brain, liver, and kidney tissues were mounted onto DNA nanoball (DNB)-patterned Stereo-seq chips. ssDNA staining and H&E staining were conducted separately for cellular localization in Stereo-seq, while DAPI staining was used for Stereo-CITE-seq; specific tissue types corresponding to each staining method are detailed in Table 2. Subsequent steps including tissue permeabilization, in situ reverse transcription, cDNA amplification, library preparation, and sequencing were performed according to the established protocol for each technology.

#### Supplementary Note 6:ssDNA Staining (Stereo-seq)

Samples for ssDNA staining were processed for Stereo-seq per the Stereo-seq Transcriptomics Set V1.3 for Chip-on-a-slide User Manual. Optimal cutting temperature (OCT)-embedded tissues were cryosectioned at 10 μm, mounted onto Stereo-seq chips, and fixed in pre-cooled methanol (−20°C) for 30 minutes. Residual liquid was removed, and chips were air-dried in a fume hood for 4–6 minutes to ensure complete methanol evaporation. Sections were stained with Qubit ssDNA Reagent (Invitrogen, Cat. No. Q10212) for 5 minutes at room temperature in the dark, after which chips were rinsed once, residual liquid was removed, and sections were mounted with Glycerol prior to image acquisition.

After imaging, permeabilization was performed using Permeabilization Mix for an optimized duration as specified in the Stereo-seq Permeabilization Set for Chip-on-a-slide User Manual.

#### Supplementary Note 7:H&E Staining (Stereo-seq)

Samples for H&E staining were processed for Stereo-seq per the Stereo-seq Transcriptomics Set V1.3 for Chip-on-a-slide User Manual. OCT-embedded tissues were cryosectioned at 10 μm, mounted onto Stereo-seq chips, and fixed in pre-cooled methanol (−20°C) for 30 minutes. Chips were stained with pre-cooled alcohol-soluble eosin (Sangon Biotech, Cat. No. A600190-0025, −20°C) for 3–5 minutes, then transferred back to the aforementioned pre-cooled methanol for an additional 1-minute fixation; residual liquid was removed thereafter.

Chips were air-dried in a fume hood for 4–6 minutes to ensure complete methanol evaporation, followed by staining with hematoxylin (Sigma, Cat. No. 51275) for 7 minutes; residual hematoxylin was decanted afterward. Chips were rinsed twice, incubated in bluing solution (Agilent, Cat. No. CS702) for 2 minutes (residual solution decanted), and rinsed once more before sections were mounted with H&E Mounting Medium for imaging. Post-imaging permeabilization was performed as described in the ssDNA staining procedure.

#### Supplementary Note 8:DAPI Staining (Stereo-CITE-seq)

DAPI staining was conducted for Stereo-CITE-seq following the Stereo-CITE Proteo-Transcriptomics Set User Manual. OCT-embedded liver tissues were cryosectioned at 10 μm and mounted onto Stereo-seq chips, which were fixed with 4% paraformaldehyde (PFA) at room temperature for 5 minutes, blocked with Blocking Buffer for 20 minutes, and incubated with TotalSeq™-A antibody mix (BioLegend, Cat. No. 199901) at room temperature for 45 minutes. Sections were washed and stained with DAPI prior to imaging.

Post-imaging, sections were washed with Wash Buffer, treated with 70% DMSO for 5 minutes to remove nonspecific antibody-oligo conjugates, rinsed with 0.1× SSC, and immediately fixed in pre-cooled methanol (−20°C) for 20 minutes. Permeabilization was performed using Permeabilization Mix for an optimized duration as detailed in the Stereo-seq Permeabilization User Manual for Stereo-CITE.

#### Supplementary Note 9:Subsequent cDNA Processing, Library Preparation, and Sequencing (ssDNA/H&E staining)

Following reverse transcription, cDNA was released from the chips using cDNA Release Mix and collected. The cDNA fraction was amplified with 4× cDNA PCR Mix and cDNA Primer, then purified via 0.8× bead-based selection (VAHTS DNA Clean Beads or AMPure® XP). Final sequencing libraries were prepared with Stereo-seq-related library reagents and sequenced on the MGI DNBSEQ-T7 or DNBSEQ-G400 platform.

#### Supplementary Note 10:Subsequent cDNA Processing, Library Preparation, and Sequencing for Stereo-CITE-seq (DAPI staining)

Following reverse transcription, cDNA and antibody-derived tags (ADTs) were released from the chips using cDNA Release Mix and collected. The two fractions were separated via 0.8× bead selection: the ADT-containing supernatant was further purified with 2.0× (0.8× + 1.2×) bead selection. cDNA and ADTs were amplified with cDNA Primer and Protein PCR Primer Mix, respectively. Sequencing libraries were prepared with the Stereo-seq 16 Barcode Library Preparation Kit and sequenced on MGI DNBSEQ-T7 or DNBSEQ-G400, enabling simultaneous profiling of the transcriptome and quantification of 128 proteins.

#### Supplementary Note 11:Processing of stereo-seq raw data

Stereoseq data were processed using SAW pipeline to obtain the expression matrix as GEF format, which includes Stereo-cell data. Briefly, The raw fastq contains coordination identity (CID) and Unique molecular identifiers (UMI) sequences.
The was aligned to reference genome (mm10 for mouse data, hg38 for human data and mRatBN7.2 for rat reference ) using STAR.

### Supplementary Figures

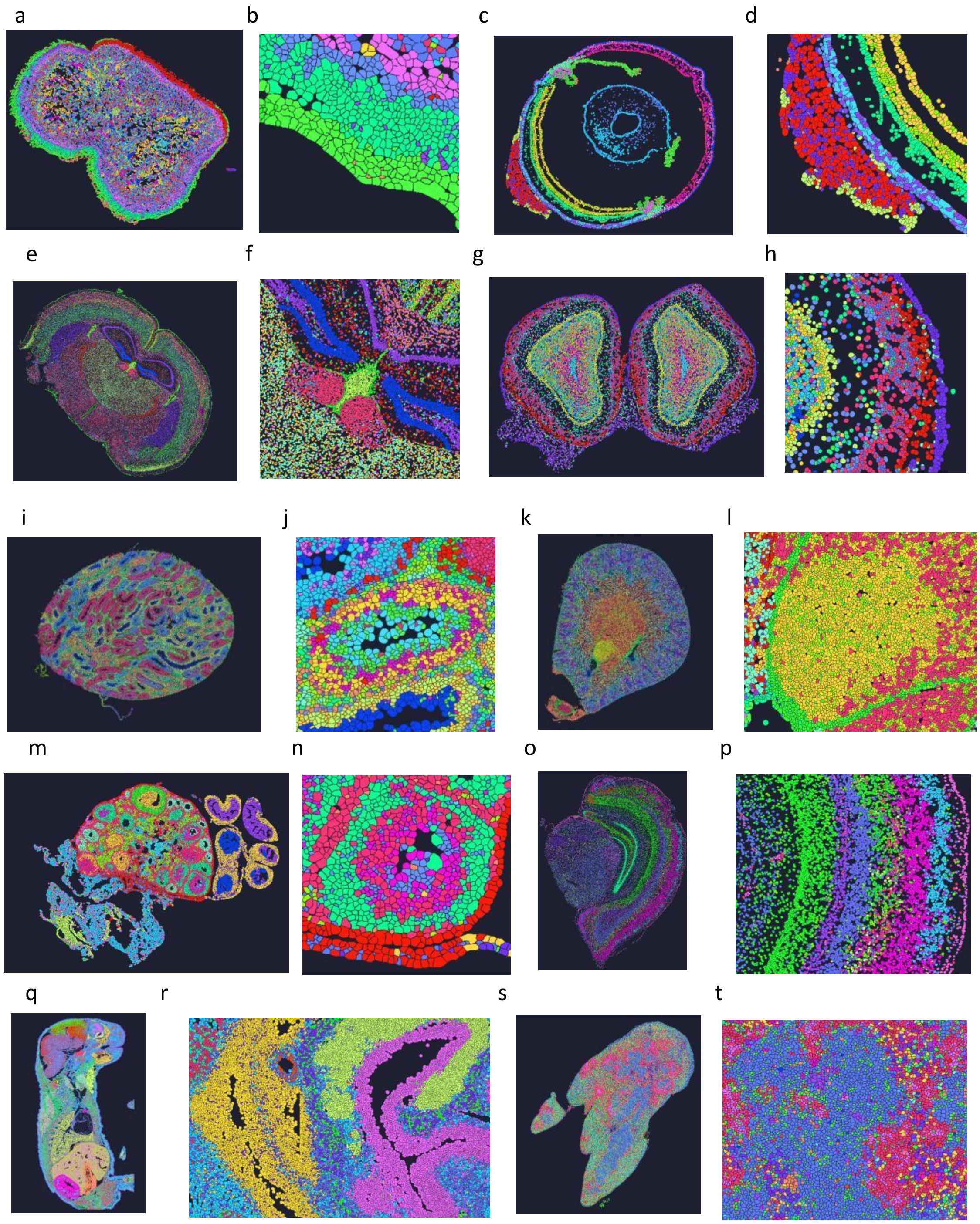

**Supplementary Figure 1.** Clustering Results of CellBin Demo Data (Download from https://en.stomics.tech/resources/demo-data/list.html or CNP0007817 of CNGBdb). Whole-mount images and their local regions: mouse tongue (whole: a; epithelium: b), eye (whole: c; lens: d), brain (whole: e; third ventricle: f), olfactory bulb (whole: g; cortex: h), testis (whole: i; seminiferous tubule: j), kidney (whole: k; inner medulla: l), ovary (whole:

m; follicle: n), brain (whole: o; cerebral cortex: p), embryo (whole: q; ocular region: r), spleen (whole: s; medulla & cortex: t).

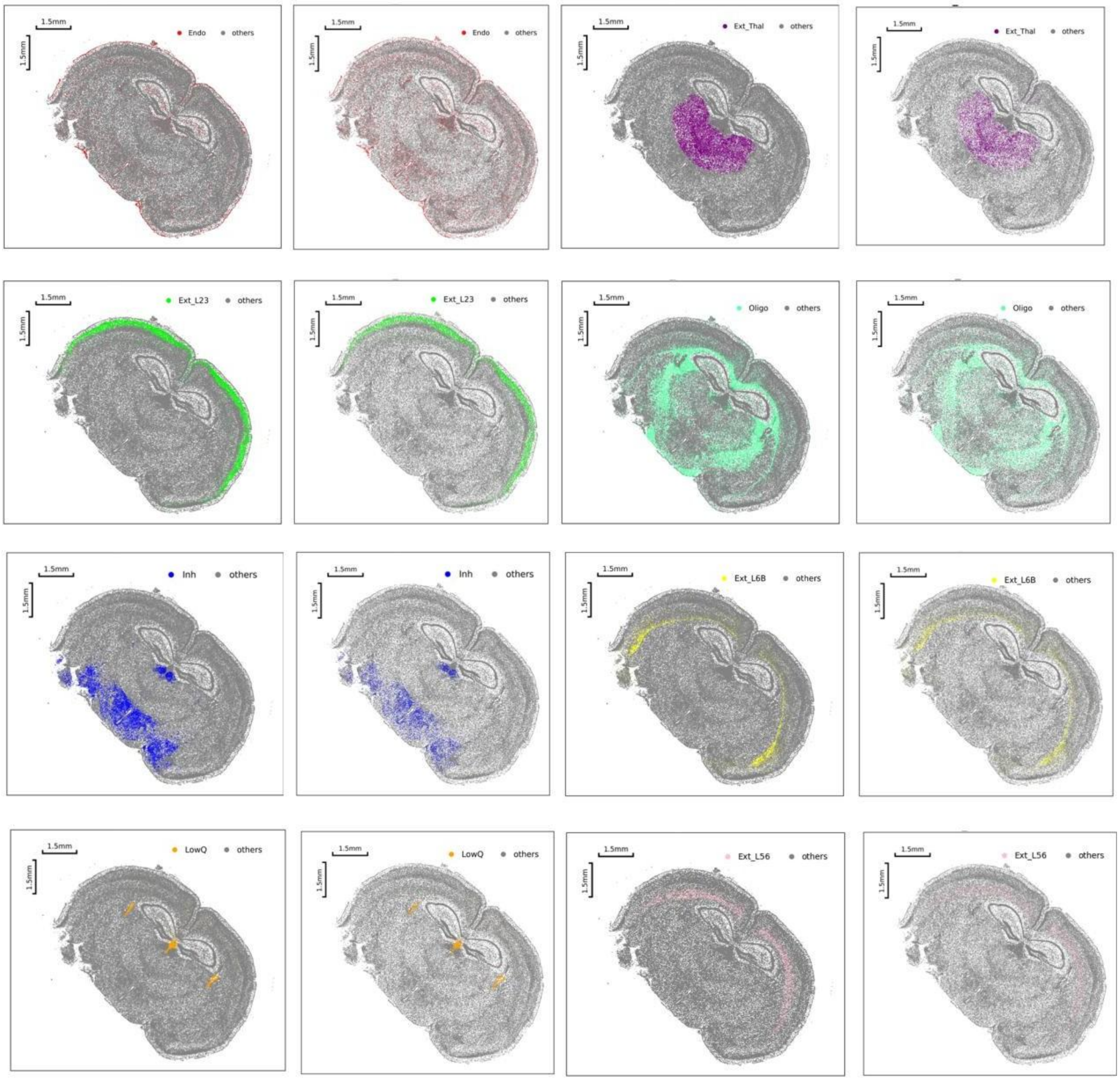

**Supplementary Figure 2:** Individual spatial distribution heatmaps of specific cell types with distinct spatial profiles from the main text-selected mouse brain dataset. Each set consists of two images: the first one is the data after cellbin segmentation, and the second one is the data after Cellpose segmentation.

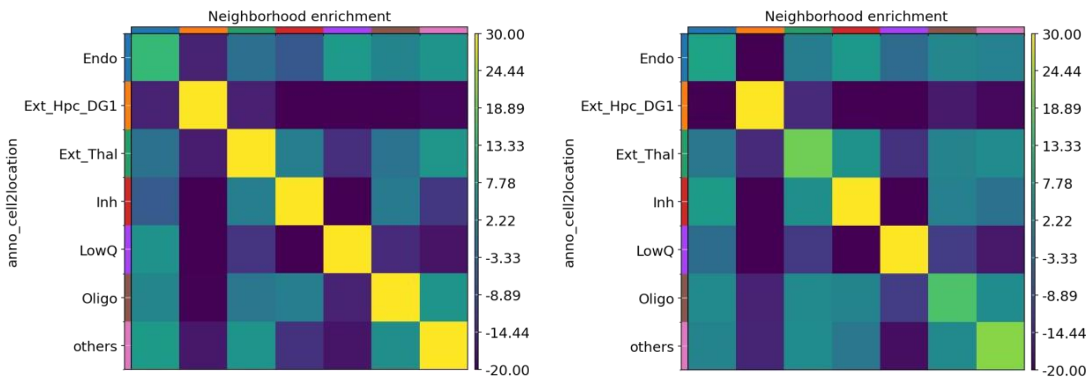

**Supplementary Figure 3:** Neighborhood Analysis of Selected Cell Types in the Habenula Region from the Mouse Brain Dataset (Cellbin on the Left, Cellpose on the Right).It can be observed that the spatial distribution of various cell types in the data segmented by Cellbin aligns more closely with the actual cell distribution in tissues—specifically, cells of the same type tend to aggregate more in a particular region, a pattern that aligns with their role in performing specialized functions.

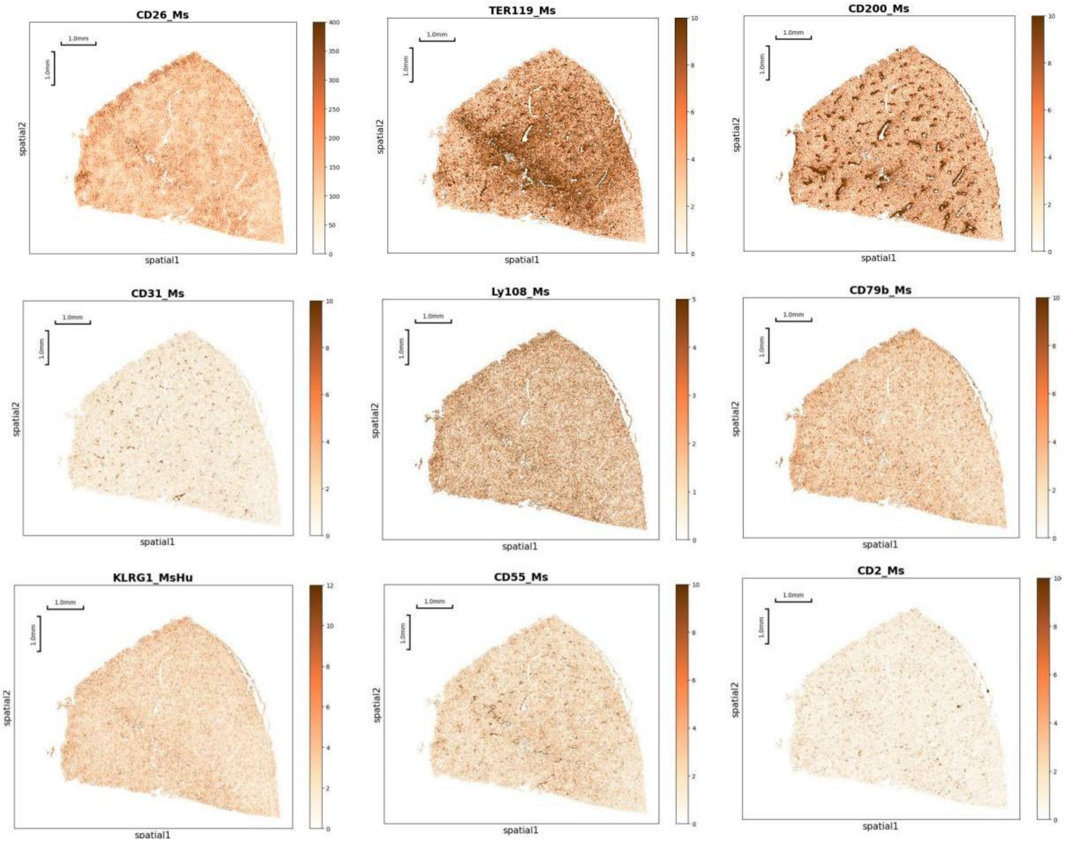

**Supplementary Figure 4:** Heatmaps of Spatial Distribution for 9 Randomly Selected Proteins in the Mouse Liver Dataset

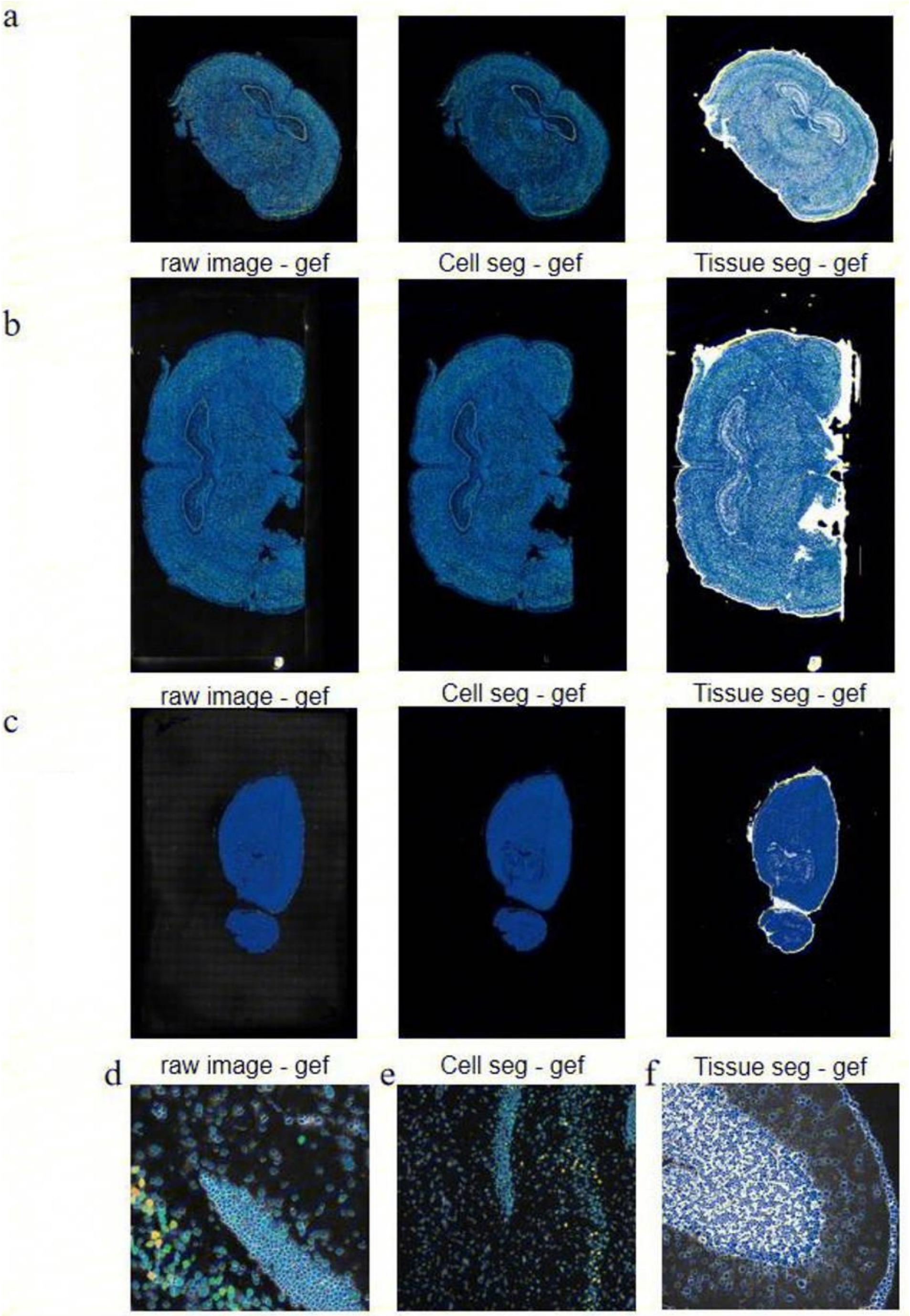

**Supplementary Figure 5:** A complete image dataset of chip samples including three key image types: raw images, cell segmentation images, and tissue segmentation images. Figures a, b, and c correspond to the three image types of samples with different sizes and IDs: 1cm×1cm (mouse brain), 1cm×2cm (rat brain), and 2cm×3cm (rat brain). Figures d, e, and f are the local magnifications of the cell segmentation images of the above three samples. Cellbin operates stably on both small and large chip samples. Moreover, its cell segmentation results clearly outline cell contours with accurate boundaries and no obvious over- or under-segmentation, further confirming reliable performance in multi-size chip scenarios.

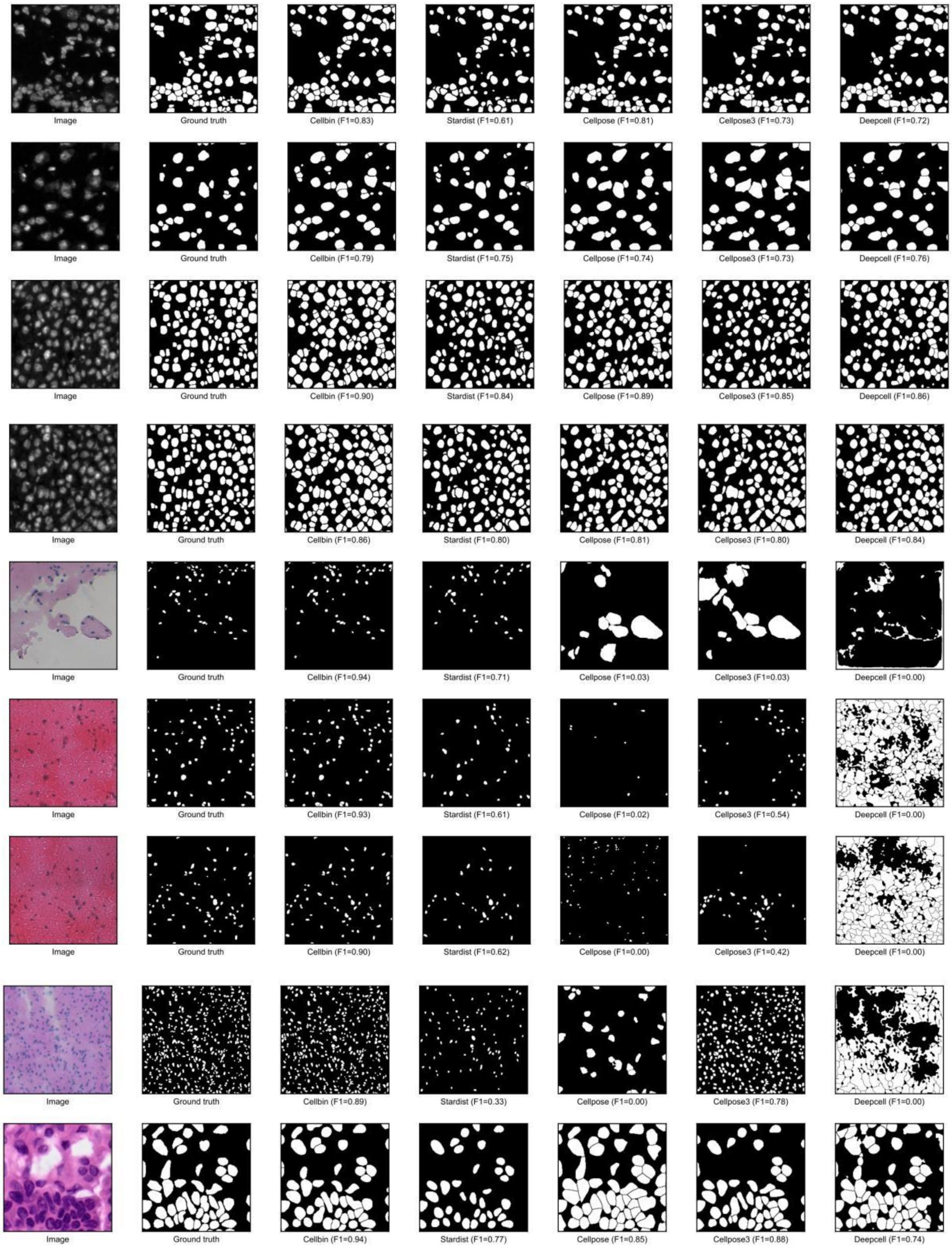

**Supplementary Figure 6:** Through the comparative demonstration of images segmented by 9 sets of softwares and the comparison of F1 scores, it can be seen that CellBin exhibits superior cell segmentation performance compared to other softwares.

### Supplementary Tables

| Technology | Resolution | Target/Capture size | Compatible Staining Methods | Throughput | Literature or links for information acquisition |
|---|---|---|---|---|---|
| 10×Genomics VisiumHD WT | 2μm | 6.5 x 6.5 mm | HE,IF | High | https://www.10xgenomics.com/platforms/visium/ product-family |
| 10×Genomics VisiumHD 3’ | 2μm | 6.5 x 6.5 mm | HE,IF | High | https://www.10xgenomics.com/platforms/visium/ product-family |
| Decoder-seq | 15μm | 5mm×5mm,7.5mm×7.5mm, 10mm×10mm.,3mm×3mm , 6mm×6mm. | HE | High | Decoder-seq enhances mRNA capture efficiency in spatial RNA sequencing |
| HDST | 2μm | Size of a single array area: The area is 13.7 mm², with specific dimensions of 5.7 mm × 2.4 mm. | HE | High | High-definition spatial transcriptomics for in situ tissue profiling |
| Slide-seqV2 | 10μm | NA | HCR-FISH, DAPI | High | Highly sensitive spatial transcriptomics at near-cellular resolution with Slide-seqV2 |
| Slide tag | 10μm | NA | Nissl Staining, HE , DAPI | High | Slide-tags enables single-nucleus barcoding for multimodal spatial genomics |
| Stereo-seq | 2μm | 5 mm×10 mm, 10 mm×10 mm, and 10 mm×20 mm, 13.2 cm×13.2 cm(max) | Nissl Staining, ISH, Nucleic Acid Dye, HE | High | Spatiotemporal transcriptomic atlas of mouse organogenesis using DNA nanoball-patterned arrays |
| Pixel-seq | 0.325μm | NA | HE, Nucleic Acid Staining, RNAscope In Situ Hybridization | High | Polony gels enable amplifiable DNA stamping and spatial transcriptomics of chronic pain |
| MERFISH | ≥ 200 nm | 1cm²,3cm² | NA | Low | Spatially resolved, highly multiplexed RNA profiling in single cells |
| seqFISH+ | 100nm | NA | DAPI | Low | Transcriptome-scale super-resolved imaging in tissues by RNA seqFISH+ |
| osmFISH | Nanoscale | 3.8 mm² | NA | Low | Spatial organization of the somatosensory cortex revealed by osmFISH |
| CosMx™ SMI | ＜100nm | NA | DAPI、FITC、TRITC、CY5 | Thousands of RNAs + Hundreds of Proteins | https://nanostring.com/products/cosmx-spatial-m olecular-imager/ https://phoenix.tsinghua.edu.cn/info/1037/1189.htm |
| Xenium | 0.2μm | 10.5 mm x 22.5 mm | HE,IF,DAPI | Up to 400 RNA transcripts | https://www.10xgenomics.com/cn/platforms/xeni um https://biocrfgz.hkust-gz.edu.cn/cn/cores/in-situ-tissue-analysis-system/ |
| Stereo-CITE | 500nm | 0.5cm×0.5cm~13cm×13cm | DAPI,ssDNA,IF,HE | 100+proteins | https://www.stomics.tech/products/Stereo-CITE https://www.stomics.tech/col243/list |
| PhenoCycler-Fu sion | 0.25μm | 18mm×35mm~630mm² | the Akoya Motif OPAL dyes,HE | 100＋biomarkers | https://www.akoyabio.com/phenocycler/instrume nt/phenocycler-fusion/ https://brcf.westlake.edu.cn/ejpt/lspt/yqsbzs/dxbywkjzxfxxt_Phenocycler_Fusion/yqjscs.htm |
| CellScape™ Precise Spatial Proteomics | 182nm | 16 mm × 40 mm | IF | NA | https://university.nanostring.com/page/cellscape https://www.bruker.com/ru/products-and-solutions/fluorescence-microscopy/spatial-omics-solutions/canopy-biosciences-cellscape.html |
| PLATO | 25μm | NA | HE,IF | Thousands of Proteins | High-resolution spatially resolved proteomics of complex tissues based on microfluidics and transfer learning |

| | | | | | |
|---|---|---|---|---|---|
| Spatial-DMT | 10μm | NA | NA | whole-genome DNA methylation and transcriptome | Spatial joint profiling of DNA methylome and transcriptome in tissues |
| Spatial-CUT&Tag | 20μm | NA | DAPI | spatial histone modification at the whole-genome scale | Spatial-CUT&Tag: Spatially resolved chromatin modification profiling at the cellular level |
| Spatial-CITE-seq | 25μm | NA | ADTs | High-throughput (approximately 200–300) proteins and transcriptome | High-plex protein and whole transcriptome co-mapping at cellular resolution with spatial CITE-seq |
| DBiT-seq | 10 μm | NA | ADTS,IF | NA | High-Spatial-Resolution Multi-Omics Sequencing via Deterministic Barcoding in Tissue |
| STARmap | Near-Single-Cel l Resolution | NA | Nissls,IF | About 1,000 cells and 160–1,020 genes | Three-dimensional intact-tissue sequencing of single-cell transcriptional states |
| OligoFISSEQ | Near-Single-Cel l Resolution | NA | IF，DAPI | Thundreds of cells and 8,000+ genes | 3D mapping and accelerated super-resolution imaging of the human genome using in situ sequencing |
| AFADESI-MSI | 100um*100um,40um *40um,20um*20um | NA | NA | 1500+ Metabolites | A Sensitive and Wide Coverage Ambient Mass Spectrometry Imaging Method for Functional Metabolites Based Molecular Histology |
| DESI-MSI | 50μm | NA | HE,IHC | NA | Desorption electrospray ionization mass spectrometry imaging in discovery and development of novel therapies |
| MALDI-MSI | 5-50μm | NA | NA | NA | Spatial Metabolomics of the Human Kidney using MALDI Trapped Ion Mobility Imaging Mass Spectrometry |

**Supplementary Table 1**: Related Information on Multiple Spatial Omics Technologys

| Cell type | Cellbin Seg. | | Cellpose Seg. | |
|---|---|---|---|---|
| | Counts | Proportion | Counts | Proportion |
| Oligo | 31391 | 22.59% | 18591 | 21.27% |
| Ext_Thal | 14089 | 10.14% | 7745 | 8.86% |
| Inh | 11970 | 8.61% | 4981 | 5.70% |
| Inh_Meis 2 | 8505 | 6.12% | 5288 | 6.05% |
| Ext_L23 | 8290 | 5.96% | 4977 | 5.69% |
| Endo | 6413 | 4.61% | 11130 | 12.73% |
| Ext_L25 | 6337 | 4.55% | 3456 | 3.95% |
| Ext_L56 | 4994 | 3.59% | 2767 | 3.16% |
| Inh_Pval b | 4619 | 3.32% | 3135 | 3.58% |
| Ext_Pir | 4070 | 2.91% | 2253 | 2.57% |

**Supplementary Table 2:** Counts and Proportion of Cell Types in the Mouse Brain Dataset

| Cell type | Cellbin Seg. | | Cellpose Seg. | |
|---|---|---|---|---|
| | Counts | Proportion | Counts | Proportion |
| Ext_L25 | 1614 | 25.89% | 918 | 20.07% |
| Ext_L56 | 866 | 13.89% | 456 | 9.97% |
| Ext_L23 | 803 | 12.88% | 470 | 10.28% |
| Oligo | 693 | 11.12% | 500 | 10.93% |
| Inh_Pvalb | 289 | 4.64% | 179 | 3.91% |
| Ext_L5 | 272 | 4.36% | 190 | 4.15% |
| Ext_ClauPyr | 210 | 3.37% | 146 | 3.19% |
| Endo | 168 | 2.70% | 452 | 9.88% |
| Micro | 128 | 2.05% | 202 | 4.22% |

**Supplementary Table 3：** Counts and Proportion of Cell Types in the Habenula Region from the Mouse Brain Dataset.

| Model name | Preprocessing | Postprocessing |
|---|---|---|
| tissue-ssDNA stain | • Histogram-based thresholding to establish an intensity cutoff, classifying suprathreshold pixels as valid tissue regions;<br>• Resizing to 1024×4028 pixels using bicubic interpolation to meet BDC U-Net input; | • Histogram-based thresholding to establish an intensity cutoff, classifying suprathreshold pixels as valid tissue regions;<br>• Resizing to 1024×4028 pixels using bicubic interpolation to meet BDC U-Net input; |
| tissue-H&E | Dynamic range compression and percentile-based normalization to preserve critical histopathological features; | • Pixels >0 are considered foreground;<br>• Resizing back to the original resolution. |
| nuclei-ssDNA | • Nonlinear attenuation of overexposed regions to mitigate saturation artifacts;<br>• Tile-based CLAHE enhancement (kernel=128, clip limit=2.56) for local contrast optimization;<br>• Min-max normalization to [0,1] range; | • Local maxima detection to extract seed points;<br>• Watershed segmentation using detected seed points as markers for instance segmentation;<br>• Morphological refinements including boundary erosion, small-object removal, and hole filling;<br>• Conversion from instance to semantic segmentation, producing a binary mask. |
| nuclei-H&E | • Color-preserving workflow: transformed images to Lab color space;<br>• Applied CLAHE exclusively to the luminance channel (L→CL) while maintaining original a/b chromaticity components;<br>• Reconstructed enhanced RGB images (CLab→RGB) before final normalization; | • Same postprocessing steps as nuclei-ssDNA. |

**Supplementary Table 4：** Preprocessing and postprocessing pipelines for different models.

| Tissue Smape | Techniques | Stain Type | Field size | Show In Figure | Data available |
|---|---|---|---|---|---|
| Mouse Brain | Stereo-seq | **ssDNA** | 1cm × 1cm | Figure 2e-w | CNP0007817(CNGBdb) |
| Mouse Live | Stereo-CITE | **DAPI** | 1cm × 1cm | Fig 1f-i and Supplement Fig 3 | CNP0007817(CNGBdb) |
| Mouse Kindey | Stereo-seq | H&E | 1cm × 1cm | Supplement Fig ?? | CNP0007817(CNGBdb) |
| Rat Brain | Stereo-seq | ssDNA | 1cm × 2cm | Supplement Fig 5 | CNP0007817(CNGBdb) |
| Rat Brain | Stereo-seq | ssDNA | 2cm × 3cm | Supplement Fig 5 | CNP0007817(CNGBdb) |
| Rat Brain | / | ssDNA | 2cm × 2cm | Supplement Fig 3 | CNP0007817(CNGBdb) |
| PBMC | Stereo-cell | / | 3cm × 3cm | Fig 1e and Supplement Fig 4 | [17] |
| Human Lung cancer | Xenium | DAPI | 1.09cm × 0.36cm | Fig 1c | 10× Genomics website |
| Mouse Lung | Visium HD | H&E | 0.65cm × 0.65cm | Fig 1b | 10× Genomics website |
| Mouse Live | Stereo-seq | DAPI and mIF | 1cm × 1cm | Fig 1d | [23] |
| Arabidopsis Seed | Stereo-seq | DAPI and CFW | 1cm × 1cm | Fig 1d | [23] |

**Supplementary Table 5：**Datasets used in this study
